# Prevalence and preferred niche of small eukaryotes with mixotrophic potentials in the global ocean

**DOI:** 10.1101/2024.02.17.580790

**Authors:** Kaiyi Dong, Ying Wang, Wenjing Zhang, Qian Li

## Abstract

Unicellular eukaryotes that are capable of phago-mixotrophy in the ocean compete for inorganic nutrients and light with autotrophs, and for bacterial prey with heterotrophs. In this study, we ask what are the overall prevalence of eukaryotic mixotrophs in vast open oceans, and how the availability of inorganic nutrients, light, and prey affects their relative success. We utilized the *Tara Oceans* eukaryotic 18S rRNA gene and environmental context variables dataset to conduct large-scale field analysis. We also performed isolate-based culture experiments to verify growth and nutritional resource relationships for representative mixotrophic taxa. The field analysis suggested that the overall prevalence of mixotrophs was negatively correlated with nutrient concentrations and positively associated with light availability. Concentrations of bacterial prey also presented somewhat correlation with mixotrophs but to a less extent. In comparison, the culture experiments demonstrated a taxa-specific relationship between mixotrophic growth and nutrition resources, i.e., the growth of one group was significantly dependent on light availability while the other group was less affected by light when they received sufficient prey. Both groups were capable of growing efficiently with low inorganic nutrients when receiving sufficient prey and light. Therefore, our field analysis and culture experiments both suggest that phago-mixotrophy for ocean eukaryotes is seemingly to be an efficient strategy to compensate nutrient deficiency but unnecessarily light limitation. This study collectively revealed a close relationship between abiotic and biotic nutritional resources and the prevalence of trophic strategies, shedding light on the importance of light and nutrients for determining the competitive success of mixotrophs versus autotrophic and heterotrophic eukaryotes in the ocean.

## Introduction

Pico- and nano-sized small eukaryotes (<3-20 µm) represent the dominant microbial communities in ocean food webs. Through metabolic activities such as photosynthesis and phagocytosis, they regulate the flux of elements and energy between primary producers and consumers. Traditionally, all protists with inherent chloroplasts were considered strict autotrophs (only photosynthesize), while those without inherited chloroplasts were regarded as strict heterotrophs (only phagocytose). It has become increasingly evident that many of those autotrophs/heterotrophs are in fact mixotrophs, phagotrophic mixotrophs particularly, they are capable of both photosynthesis and phagocytosis [1,2,3]. Depending on whether the organism possesses its own chloroplasts, mixotrophs can be further categorized into constitutive mixotrophs and non-constitutive mixotrophs. Constitutive mixotrophs, the focus of the current study, are normally small-size flagellated eukaryotes that primarily feeding on bacterial prey [4,5], whereas non-constitutive mixotrophs are larger-sized microplankton (20-200 µm) which graze on both eukaryotic and bacterial prey [6,7]. Due to the distinct metabolic preference and trophic activities, those two mixotrophic groups should be studied separately.

The relative abundance and prevalence for mixotrophs (comparing to other eukaryotes) in a given system can be partially interpreted as the competition results among mixotrophs, autotrophs, and heterotrophs. Specifically, mixotrophs and autotrophs compete for inorganic nutrients (hereinafter referred to as nutrients) and light for autotrophic growth, and for bacterial prey (including heterotrophic and autotrophic bacteria) with bacterivorous heterotrophs for heterotrophic growth. Although results may vary, previous studies have suggested that the physiological performance and competitive advantage of mixotrophs can be related to these three nutritional resources of nutrients, light and prey [8,9,10,11]. Mixotrophy are most likely benefited from scenarios where nutrition is not sufficient to support a single trophic strategy, i.e., strict autotroph or heterotroph. For example, Ward (2019) has proposed a conceptual model arguing that mixotrophy can effectively alleviate dissolved inorganic nitrogen deficiency in oligotrophic environments via phagocytosis of high-nitrogen bacterial prey [12]. Meanwhile, it is necessary to acknowledge that the photosynthesis and/or phagocytosis performance alone is very likely penalized for mixotrophs [13], and these tradeoffs can differentiate significantly among different species. Li et al. (2022) have demonstrated a distinct variation for grazing/heterotrophic capability and trophic efficiency among different mixotrophic strains [5].

In regional areas of temperate sea [14,15,16] and oligotrophic ocean [11,17,18] or lakes [19,20,21], field observations have also shown that mixotrophy can be advantageous for phytoplankton community growth when nutrition are not ideal for autotrophy, which are mostly due to limiting nutrients and/or irradiance. In other words, conditions that are unfavorable for strict autotrophy or heterotrophy could possibly select for mixotrophy on the entire community level. Yet for larger spatial scales that encompass multiple ocean regions, there has been few studies to look into the broad-scale relationship between mixotrophic prevalence and environmental conditions related to nutrition availability. While acknowledging these relationships can be complex due to multi-factorial interactions among biotic and abiotic components in the food web, it is reasonable to speculate nutrients, light and prey can play an important role determining the relative success of mixotrophy. With climate change, evidence has shown that mixotrophs may play a more important role due to their metabolic plasticity, i.e., easier to adapt to changing environments [22,23,24,25]. Motivated by these findings, we are particularly interested to delineate the overall prevalence and realized niche favoring mixotrophs in comparison to strict autotrophs and heterotrophs in vast open oceans across the globe.

The broad goal of this study is to gain insights into the interrelationship between environmental conditions and trophic strategies on both community and species level, as well as the underlying mechanism behind it. We aim to answer the following scientific questions, 1) What are the overall prevalence of mixotrophic eukaryotes in open oceans? 2) Are there specific environments under which mixotrophs are favored over autotrophs and heterotrophs; 3) If so, what role does nutrients, light and bacterial prey play for determining the relative success and fitness of mixotrophy? We use the eukaryotic 18S rRNA gene dataset from *Tara Oceans Expedition* to assign trophic mode to each lineage/species and assess the relationship between trophic mode composition and contextual variables. Subsequently, we run culture experiments to confirm relationships between representative mixotrophic isolates and key nutritional resource variables. The *Tara Oceans* analysis revealed statistically important variables that are affiliated with the trophic mode prevalence, and the experimental results provided detailed and physiological evidence for it.

## Materials and Methods

### Study regions

The data generated by *Tara Oceans* expedition between 2009 and 2013 are particularly suited to study the genetic and functional diversity of plankton. In this study, a total of 104 stations and 165 samples were investigated, among which all 104 sites had surface samples and 61 sites had both surface and chlorophyll maximum layer (CML) samples. A surface dataset (SUR) consisting of 104 samples was used, while the surface and chlorophyll maximum depth dataset (SUR&CML) comprising 122 (the 61 stations have both SUR&CML samples) and 165 samples (all 104 SUR samples and 61 CML sample), was generated for different statistical analysis. These samples were collected during four different seasons (Spring, Summer, Autumn, Winter) between 2009 and 2013 from eight oceanic regions (Northern Atlantic Ocean, Mediterranean Sea, Red Sea, Indian Ocean, Southern Atlantic Ocean, Southern Ocean, Southern Pacific Ocean and Northern Pacific Ocean) with broad latitudinal (64.4°S-43.7°N) and longitudinal ranges (159.0°W-73.9°E). Bathymetry depths varied from 13 to 5964 km and distance to coast was between 0.9-1404 km.

### Contextual environmental variables

We built an environmental context dataset for further statistical analysis following these three steps. First, representative environmental variables were obtained from both the Tara Oceans PANGAEA data repository (http://www.pangaea.de) and a variety of publications, including Pesant et al. (2015), Picheral et al. (2017), Ardyna et al. (2017), Ibarbalz et al. (2019), Faure et al. (2019), Karlusich et al. (2022) etc. (**Supplementary Dataset S1**) [26,27,28,29,30,31]. The major approaches for retrieving these variables included in situ sensors, laboratory analysis of water samples, climatological instruments and model simulations, following standard protocols and analytical methods (details are described in those aforementioned literatures). Second, these ∼100 variables were manually filtered down to 77 by eliminating similar variables and less relevant ones, to reduce collinearity and inflation of dimensionality. Examples include salinity was chosen over conductivity, and excluding factors such as moon phase durations and current residual time etc. Lastly, when variables have values from multiple measurements and/or calibration/calculation methods, only the most accurate one or suitable one (for the purpose of this study) was selected. For example, median values for photosynthetically available radiation (PAR), averaged on an 8-day interval and calculated based on attenuation coefficient and surface irradiance observed by AMODIS were selected over a 1-day interval to avoid fluctuations caused by short-term weather conditions. Total chlorophyll *a* concentrations, derived from sensor measurements and calibrated with non-photosynthetic quenching and discrete water sample measurements, were used among other available measurements. Additional comparisons were also conducted to confirm there were no overall significant differences between the selected one and other measurements for the same variable, such as 8-day averaged PAR vs 30-day averaged PAR, total Chl *a* derived from sensor vs water samples vs MODIS, and nutrient concentrations derived from water samples and model simulation, etc.

The final contextual environmental dataset comprised 29 variables (**Supplementary Dataset S2)**, representing physical conditions such as temperature, salinity, density, attenuation coefficient of PAR (Kd PAR), chlorophyll maximum layer (CML), mixed layer depth (MLD), nitracline depth and euphotic depth; climatology such as PAR and sunshine duration; biochemical conditions such as total Chl *a*, oxygen, nutrients (NO_3_^-^&NO_2_^-^, NO_2_^-^, PO_4_^3-^, SiO_4_^-^), pH, total alkalinity, total carbon, fluorescence, particulate organic carbon (POC), particulate inorganic carbon (PIC), and bacterial abundances including autotrophic (*Prochlorococcus* and *Synechococcus*) and heterotrophic bacteria. Note that longitudes, latitudes, distance to coasts, sampling depths and bathymetry depths were not included in the analysis to better focus on the aforementioned climatological, biochemical and physical variables.

### 18S rRNA gene abundances and corrected cell abundances

Detailed genomic sample processing methods were described in de Vargas et al. (2015) [32]. Briefly, community genomic DNA was extracted from cells filtered through membrane filters with different pore sizes (0.8µm, 5µm, 20µm) after pre-filtration with nylon sieves (to retrieve different plankton communities). In this study, we focused on the 0.8-5µm size range, which contributed the majority of eukaryotic community in sampled regions [32]. The eukaryotic V9-18S rRNA gene region was sequenced through Illumina HiSeq, assembled, filtered and classified with SWARM for operational taxonomic units (OTUs). The sequence files were downloaded from the European Nucleotide Archive under project number PRJEB6610. Subsequently, we re-assigned OTUs to the lowest taxonomic rank possible, referring to the Protist Ribosomal Reference (PR^2^) database [33]. Ciliophora, Radiolaria, and Foraminifera that were likely derived from large-sized populations (contributing a small portion of the total species), were excluded from further analysis to focus on small-sized constitutive mixotrophs. Unkown Eukaryota and a small portion of parasites (e.g., Fungi, Amoebozoa, Syndiniales, Dinoflagellata) were excluded from further trophic annotation, and thus further analysis. This final gene dataset has generated 40,126,831 sequences and 845 lineages/species for surface samples, and 17,874,753 sequences and 803 lineages/species for the CML samples, respectively (**Supplementary dataset S3**).

Eukaryotic communities are known to exhibit substantially different 18S rRNA gene copy numbers, varying from as low as one copy in species belonging to Ochrophyta to thousands of copies per cell in dinoflagellates [5,34]. Overall, cellular biovolumes have been shown to be positively correlated with 18S rRNA gene copy numbers but certain taxa, such as dinoflagellates, tended to have particularly high copy numbers [35,36]. To correct for this bias and obtain gene-converted-cell abundances, we first collected empirical data from plankton, focusing on small cells with equivalent diameter of 5-10 µm and biovolume of 65-523 µm^3^ (**Supplementary Table S1**). Then, we fit a linear correlation equation between empirical 18S rRNA gene copy numbers and cellular biovolume for individual group of dinoflagellates, diatoms, and other eukaryotes (including chlorophytes, haptophytes, dictyochophytes, etc.), respectively. Finally, a correction factor (C.F.) for each group was calculated using upper limit biovolume of 65 µm^3^, that gives 59.3, 10.5, and 4.7 for dinoflagellates, diatoms, and other eukaryotes, respectively. For comparison, Martin et al. (2022) used 27.1, 4.4, and 0.9 C.F. for dinoflagellates, diatoms, and all other eukaryotes [36]. Gene abundances were converted into cell abundances by dividing gene numbers with C.F. for each group. The corrected cell abundance was the focus for further analysis, but 18S rRNA gene abundances were also assessed occasionally for comparison purposes.

### Assignment of trophic mode

In this study, we focus on the trophic strategy potentials among different eukaryotic lineages, rather than the in situ active trophic activities (are partially associated with instantaneous conditions). Therefore, we assigned each lineage/species (when possible) into three trophic groups by searching published database and references [5,37,38], including (potential) mixotrophs (pigmented eukaryotes with phagocytosis capability), heterotrophs (eukaryotes with no inherent chloroplasts) and autotrophs (pigmented eukaryotes with no available evidence for phagocytosis). When trophic annotation was impossible at lineage or species level, their affiliated genus or family was referred to for annotation. The filtration has generated a final composition of 184-186 mixotrophic, 283-313 heterotrophic, and 303-377 autotrophic lineages/species for the SUR and SUR&CML samples, respectively (**Supplementary dataset S3**). We acknowledge the limitation of this trophic annotation methodology, and address it further in the Discussion.

### Trophic index and trophic composition

The relative prevalence or success of each trophic group at a given site was expressed as the trophic index of each group (TI*_g_*), where *g* represents three trophic groups of mixotroph (M), autotroph (A) and heterotroph (H). Therefore, TI*_g_* of TI_M_, TI_A_, and TI_H_ can be quantified as their relative abundance against total eukaryotic community (T), either in cell or gene abundances (expressed as M:T, A:T and H:T). The combined three index of TI_M_, TI_A_ and TI_H_ can represent the complete results for trophic mode compositions at a given site.

### Statistical analysis

The surface dataset was used for redundancy analysis (RDA) to elucidate the overall distribution pattern of trophic groups among 104 stations, and potential relationships with environmental variables. All 29 filtered contextual variables were considered for the analysis, with missing values for each variable replaced with means and then log-transformed for standardization prior to analysis. Hellinger transformation was applied to species abundances and escoufier selection with a threshold of 0.9 was applied to the trophic species. Environmental variables that passed significance test and collinearity test were presented on the RDA plot. The RDA was carried out in R with the vegan package, following the script outlined by Faure et al. (2019) [30]. Principal components analysis (PCA) was applied to extract the major components that explained the environmental gradients from both surface (SUR) and SUR&CML datasets with R package pcaMethods [39]. After extracting the main variable components, TI*_g_*indices were plotted over two axes of PC1 and PC2 with generalized additive model. Stepwise regression between all filtered variables and TI_M_ were conducted to retrieve important variables explaining mixotrophic prevalence, and linear regression between single variables and TI*_g_* were carried out to provide additional correlation results. ANOVA was conducted for the culture experiments to reveal factorial/treatment impact on growth rates and grazing rates among various mixotrophic isolates.

### Culture experiments

Methods for isolating and cultivating mixotrophs from the North Pacific Subtropical Gyre were described in Li et al. (2021; 2022) [5,40]. Briefly, mixotrophic eukaryotes were isolated from the euphotic zone at Station ALOHA (22° 45’ N, 158° 00’ W) by enriching seawater samples in nitrogen-depleted Keller medium (K-N medium) that supplemented with bacterial prey. Six identified isolates belonging to Dictyochophyceae, Prymnesiophyceae, and Chrysophyceae were selected for conducting growth experiments. These strains were phylogenetically close to the *Tara Oceans* lineages that were among the top 15 abundant lineages. Prior to experiments, all isolates were acclimated for 1-2 months (duplicated over 10 generations) under corresponding treatment conditions, i.e., low nutrients (K-N medium) and high light, low nutrients and low light, high nutrients (K medium) and high light, and high nutrients and low light. The high and low light treatments were receiving ∼100 and ∼10 µM photons m^-2^ s^-1^ irradiance on a light/dark cycle (12/12 h). All treatments were supplemented with autotrophic bacterial prey of *Prochlorococcus* (strain MIT9301) at final concentrations of ∼2×10^6^ cells mL^-1^, which represents the second abundant prokaryotic population at station ALOHA. Prey cells were centrifuged with a rotation speed of 2000-3000 × g for 3–5 min and gently resuspended for the enrichment. Controls without added prey or grazers were also set up at each condition for comparison purposes. Cell concentrations of prey and grazers were measured every 12–24 hours by flow cytometry, and ingestion evidences were occasionally obtained via microscopic observation.

Based on our previous study on the grazing functional responses, most isolates have ingestion rates saturated when prey concentration are higher than 10^6^ cells mL^-1^. Meanwhile, clearance rates (ingestion rates divided by prey concentration) can roughly present the maximal clearance rates with the final prey concentration used in these experiments. Therefore, we use clearance rates instead of ingestion rates [1][41] to present grazing performance. Ingestion rates (prey grazer^-1^ h^-1^) for each grazer were calculated as the amount of prey removed between sampling time t+1 and t, divided by averaged grazer concentrations over the same time period. Clearance rates were calculated by dividing the ingestion rate by averaged prey concentration over the same interval. Body volume specific clearance rates (body volume grazer^-1^ h^-1^) were calculated by dividing the clearance rate with cellular biovolume (µm^3^) of each grazer (measured under microscopy) [42]. Clearance rates reported in the results were averaged over all sampling intervals prior to grazers reaching stationary phase. Growth rates presented in the results were maximal values during exponential growth phase (slope of Ln transformed cell abundance versus time).

## Results

### Oceanographic characteristics across the *Tara Oceans* sampling regions

The 104 surface samples encompassed eight ocean habitats on large spatial scale (**Fig. 1a**). Geographically they span roughly four latitudinal regions, from low (0°-20°) to low-medium (20°-30°), to medium (30°-40°), and medium-high latitude (40°-60°). Moving from low to high latitudinal regions, overall PAR decreased from 25.4 to 14.7 mol photons m^-2^ d^-1^, nutrients, e.g., NO_3_^-^&NO_2_^-^, increased from 0.03 to 17.3 µmol L^-1^, and heterotrophic bacterial abundances were relatively stable, ranging between 3.9-5.3 ×10^5^ cells mL^-1^ (**Table 1**; **Fig. 1b**). The globally averaged means were 21.3 mol photons m^-2^ d^-1^ of PAR, 0.14 µmol L^-1^ of NO_3_^-^&NO_2_^-^ and 4.5 ×10^5^ cells mL^-1^ of heterotrophic bacteria, demonstrating most sampling sites were oligotrophic environments. Smaller regional variations were also observed. The higher nutrient concentrations in low latitude regions were possibly due to the Peruvian coastal and eastern equatorial upwelling, while lower PAR in medium latitude areas was because Mediterranean Seas were sampled during winter.

**Fig. 1.**
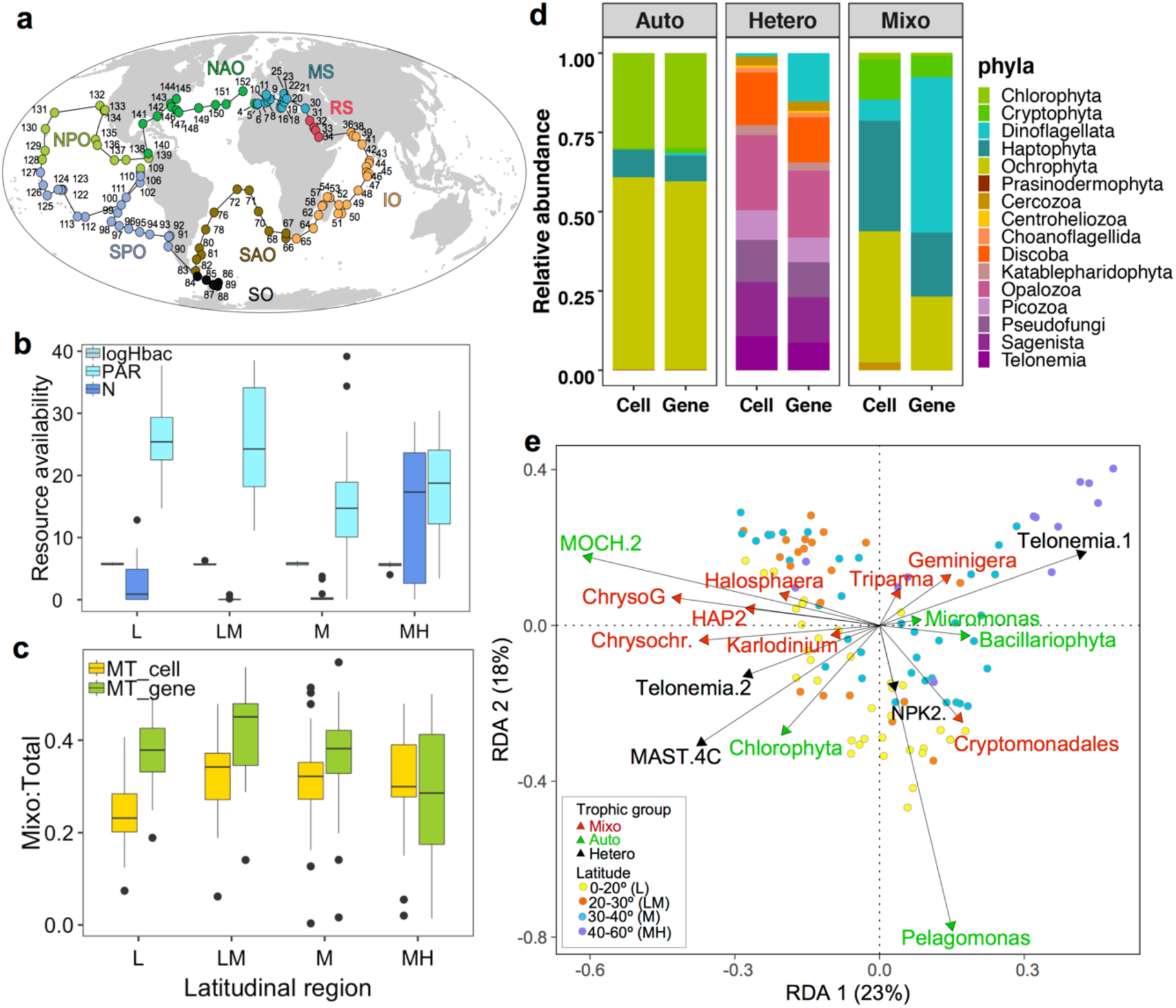
Map of study sites (**a**), averaged median values of inorganic nutrients (NO_3_^-^&NO_2_^-^; N for abbreviation), light (PAR) and heterotrophic bacteria (**b**), and mixotrophic prevalence index TI_M_ (Mixo:Total) (**c**) at different latitudinal regions. Panel **d** presents composition of autotrophic (Auto), mixotrophic (Mixo) and Heterotrophic (Hetero) eukaryotes on phyla level derived from 18S rRNA gene and corrected cell abundances. Panel **e** is the redundancy analysis plot with distribution of trophic groups on species level. Selected RDA variables and *Tara Oceans* lineage full names can be found in Supplementary Fig. S1. Latitudinal region name and definition were shown in the legend of panel **e**.

**Table 1.** Oceanographic characters and trophic index of surface water samples. Abbreviations for PAR, NO_3_^-^&NO_2_^-^, Hbac stands for photosynthetically available radiation, dissolved nitrate and nitrite (NO_3_^-^&NO_2_^-^), and heterotrophic bacteria, respectively. M:T, M:A and M:H denotes rations of either cell abundance (cell) or 18S rRNA gene abundance (gene) among mixotroph (M), autotroph (A), heterotroph (H) and total eukaryotes (T).

| Latitude | Oceanic Region<sup>†</sup> | Station | Temp (°C) | Salinity | PAR (mol m<sup>-2</sup> d<sup>-1</sup>) | $\text{NO}_3^-$ & $\text{NO}_2^-$ (μmol L<sup>-1</sup>) | Hbac (cells mL<sup>-1</sup>) | M:T (Cell /Gene) | A:T (Cell /Gene) | H:T (Cell /Gene) |
| --- | --- | --- | --- | --- | --- | --- | --- | --- | --- | --- |
| 0-20° | SPO, IO, NPO, SAO, NAO | 36 | 26.6 | 35.1 | 25.4 | 0.89 | 4.9e5 | 0.23/0.38 | 0.32/0.24 | 0.43/0.38 |
| 20-30° | IO, NPO, SPO, SAO, RS | 17 | 24.7 | 36.2 | 24.3 | 0.03 | 3.9e5 | 0.34/0.45 | 0.10/0.08 | 0.51/0.44 |
| 30-40° | MS, SAO, NAO, SO | 37 | 19.5 | 36.4 | 14.7 | 0.13 | 5.3e5 | 0.32/0.38 | 0.19/0.16 | 0.48/0.43 |
| 40-60° | SO, SAO, MS | 14 | 7.1 | 34.3 | 18.8 | 17.3 | 4.5e5 | 0.30/0.29 | 0.18/0.22 | 0.47/0.46 |
| Global | SPO, IO, NPO, SAO, NAO, SO, RS, MS | 104 | 23.8 | 35.4 | 21.3 | 0.14 | 4.5e5 | 0.29/0.38 | 0.21/0.19 | 0.47/0.42 |

<sup>†</sup> Oceanic regions follow the Tara Oceans project standards. NAO: North Atlantic Ocean; MS: Mediterranean Sea; RS: Red Sea; IO: Indian Ocean; SAO: South Atlantic Ocean; SO: Southern Ocean; SPO: South Pacific Ocean; NPO: North Pacific Ocean.

### Relative abundances and taxonomic compositions of three trophic modes

The trophic index of TI_M,_ TI_A_ and TI_H_ varied from 0.23 to 0.34, 0.10 to 0.32 and 0.43 to 0.51 in cell abundance, and from 0.29 to 0.45, 0.08 to 0.24 and 0.38 to 0.46 in gene abundance, respectively (**Table 1**; **Fig. 1c**). TI_M_ in cell abundance were overall somewhat lower than gene abundance. Nevertheless, the highest TI_M_ median values were both retrieved from low-medium latitude, and the lowest values were from low latitude (cell abundances) and medium-high latitudes (gene abundances), respectively. In comparison, high and low TI_A_ median values were from low and medium-high latitude and low-medium latitude (both cell and gene abundances). TI_H_ did not change as much across latitudinal gradients, but highest median values in cell and gene abundances were shown in low-medium and medium-high latitudes, respectively.

Before the gene-cell abundances correction, mixotrophs were mostly composed of lineages from Dinoflagellata, Haptophyta and Ochrophyta. After the correction for cell abundances, the dominant mixotrophs were retrieved from Haptophyta and Ochrophyta (**Fig. 1d**). Proportions of dinoflagellates decreased for heterotrophic community after gene-cell conversion, and the rest three most abundant heterotrophs were belonging to phyla of Opalozoa, Discoba and Sagenista, both in cell and gene abundances. Differences for autotrophic compositions on phyla level in cell and gene abundances were insignificant, which were both dominated by Chlorophyta and Ochrophyta. Redundancy analysis suggested stations from the same ocean or latitudinal regions tended to cluster together (**Fig. 1e; Supplementary Fig. S1**). Certain species were more abundant in lower latitudes, such as Chrysophyceae_Clade G_sp. of Ochrophyta, *Chrysochromulina* sp. and Haptophyta_HAP2_XXX_sp. of Haptophyta, and *Karlodinium veneficum* of Dinoflagellata for mixotrophs, MOCH.2_XXX sp. and Chlorophyta_XXXX_sp. of Chlorophyta for autotrophs, and MAST.4C_XX sp. and Telonemia Goup2_X sp. for heterotrophs. Those taxa were positively affiliated with PAR and CML, and negatively correlated with nutrients (NO_3_^-^&NO_2_^-^ and NO_2_^-^) and other variables located on the right side of the RDA plot. Other species, including *Geminigera cryophila* of Cryptophyta and *Triparma mediterranean* of Ochrophyta for mixotrophs and Telonemia Group1 for heterotrophs, were abundant at higher latitudes and positively correlated with oxygen and sunshine duration. Additionally, *Micromonas* A2 and Bacillariophyta_XXX sp. for autotrophs were associated with high MLD, POC and NO_2_^-^, while Cryptomonadales_XX_sp. for mixotrophs and NPK2 lineage_X_sp. for heterotrophs were affiliated with high abundances of *Prochlorococcus*.

### Nutritional variables and the prevalence of three trophic modes

Among the 29 selected predictors, availability of nutrients (including NO_3_^-^&NO_2_^-^ and silicate), PAR and temperature were the top variables revealed by principal component analysis for explaining the environmental conditions of surface (SUR) and surface and chlorophyll maxima layer (SUR&CML) samples (**Table 2**; **Figs. 2, 3**). Specifically, both principal component axes (PC1 and PC2) for the SUR&CML samples were mostly explained by PAR and nutrients (loading coefficients varied from −0.42 to 0.17; R^2^ of 0.30). When only considering SUR samples, PC1 axis was mostly contributed by nutrients and temperature (R^2^ of 0.35), while PC2 axis (R^2^ of 0.24) were mostly explained by MLD, CML and PAR.

**Table 2.** Major environmental variables retrieved by principal component analysis for surface (SUR) and surface and chlorophyll maximal layer (SUR&CML) datasets. Top variables from each principal component axis (PC1 and PC2) for both datasets were shown with their loading coefficients (marked in bold). Symbols of ‘+’ and ‘−’ explain the direction of changing variable values from left to right on the axes.

|  | SUR |  | SUR& DCM |  | sum<sup>†</sup> |
| --- | --- | --- | --- | --- | --- |
|  | PC1 | PC2 | PC1 | PC2 |  |
| PAR | -0.08 | **-0.14** | **-0.42** | **-0.35** | 0.99 |
| NO<sub>3</sub><sup>-</sup>&NO<sub>2</sub><sup>-</sup> | **0.31** | **-0.14** | **0.23** | **-0.25** | 0.93 |
| Silicate | **0.23** | -0.10 | **0.17** | **-0.17** | 0.67 |
| Temperature | **-0.24** | 0.08 | -0.14 | 0.13 | 0.59 |
| Nitracline depth | -0.20 | -0.11 | -0.15 | 0.12 | 0.58 |
| CML | 0.06 | **-0.17** | -0.10 | 0.07 | 0.40 |
| Mixed layer depth | 0.06 | **-0.22** | -0.04 | -0.06 | 0.38 |
| R<sup>2</sup> | 0.35 | 0.24 | 0.30 | 0.25 |  |

<sup>†</sup> summed absolute values from PC1 and PC2 for both datasets

**Fig. 2.**
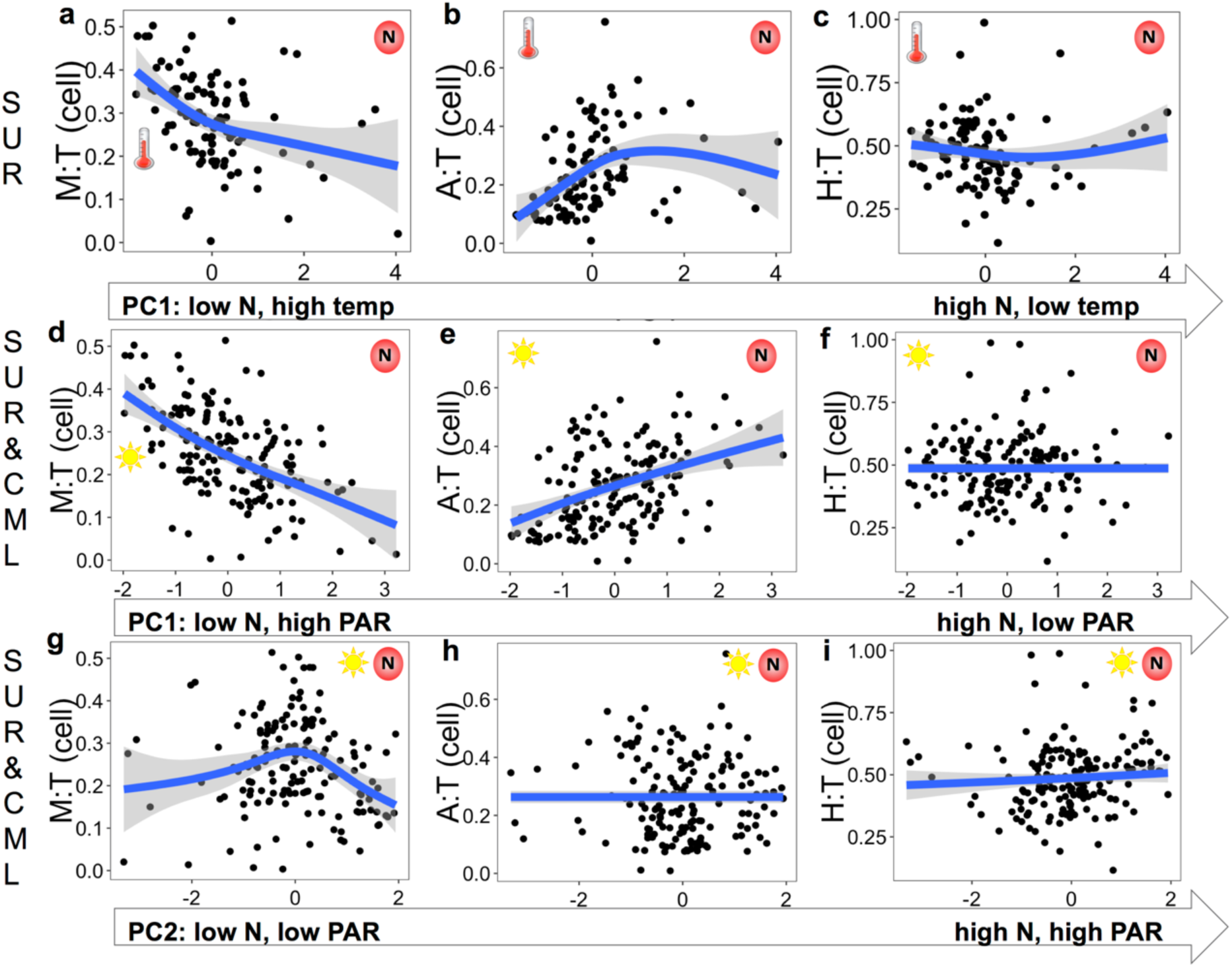
Variation patterns of TI_A/M/H_ in cell abundances along PC1 axis for surface (SUR) dataset (**a-c**), and PC1 (**d-f**) and PC2 (**g-i**) axes for surface and chlorophyll maximal layer (SUR&CML) dataset. Principal components of nutrients (NO_3_^-^&NO_2_^-^), PAR, and temperature were marked on each panel, and the location of symbols indicates higher values ends. Labels embedded on each PC-axis indicate how variables change from left to right. Blue lines and shaded bands were fitted generalized additive model smoother and 95% confidence interval.

The fitted smoothers of TI*_g_* demonstrating the most substantial responses were from mixotrophs to changing principal components. Specifically, there was a decreasing trend of TI_M_ along PC1 for both datasets, indicating decreasing prevalence of mixotroph with increasing nutrients for SUR (**Fig. 2a**), as well as decreasing PAR and increasing nutrients for SUR&CML (**Fig. 2d**). TI_M_ did not demonstrate clear changes with the PC2 axis from SUR samples so results were not shown. It is noteworthy that the slope for decreasing TI_M_ when nutrients are increasing and PAR were decreasing was the steepest among all plots (**Fig. 2d**). For PC2 axes of SUR&CML dataset, TI_M_ presented a unimodal shape with changing nutrients and PAR conditions, with maximal values obtained around zero PC2 scores (**Fig. 2g**). These results may suggest a more complex non-linear relationship between TI_M_ and co-varying nutrients and light conditions. TI_A_ overall increased with increasing nutrients for the SUR dataset (**Fig. 2b**), and increasing nutrients and decreasing PAR for the SUR&CML dataset (**Fig. 2e**). When nutrients and PAR changed in the same direction (PC2 for SUR&CML dataset), TI_A_ did not show clear variations (**Fig. 2h**). TI_H_ overall varied much less with changing conditions compared to TI_M_ and TI_A_ (**Figs. 2c, f, i**), among which a weak unimodal shape was observed with PC1 from SUR dataset (**Fig. 2c**). For the gene abundances denoted results, similar trends were found for TI*_g_* variations over the principal components but with different slope shapes (**Fig. 3)**. Particularly, a steeper decreasing slope was presented for TI_M_ on PC1of SUR samples and a steeper increasing slope (before the peak) was seen on PC2 for the SUR&CML samples (**Fig. 3a, g**). The weak increasing trend of TI_H_ over PC1 axis of SUR samples disappeared (**Fig. 3c)**, and a somewhat stronger increase of TI_H_ were demonstrated along both axes of PC1 and PC2 for the SUR&CML dataset (**Fig. 3f, i).**

**Fig. 3.**
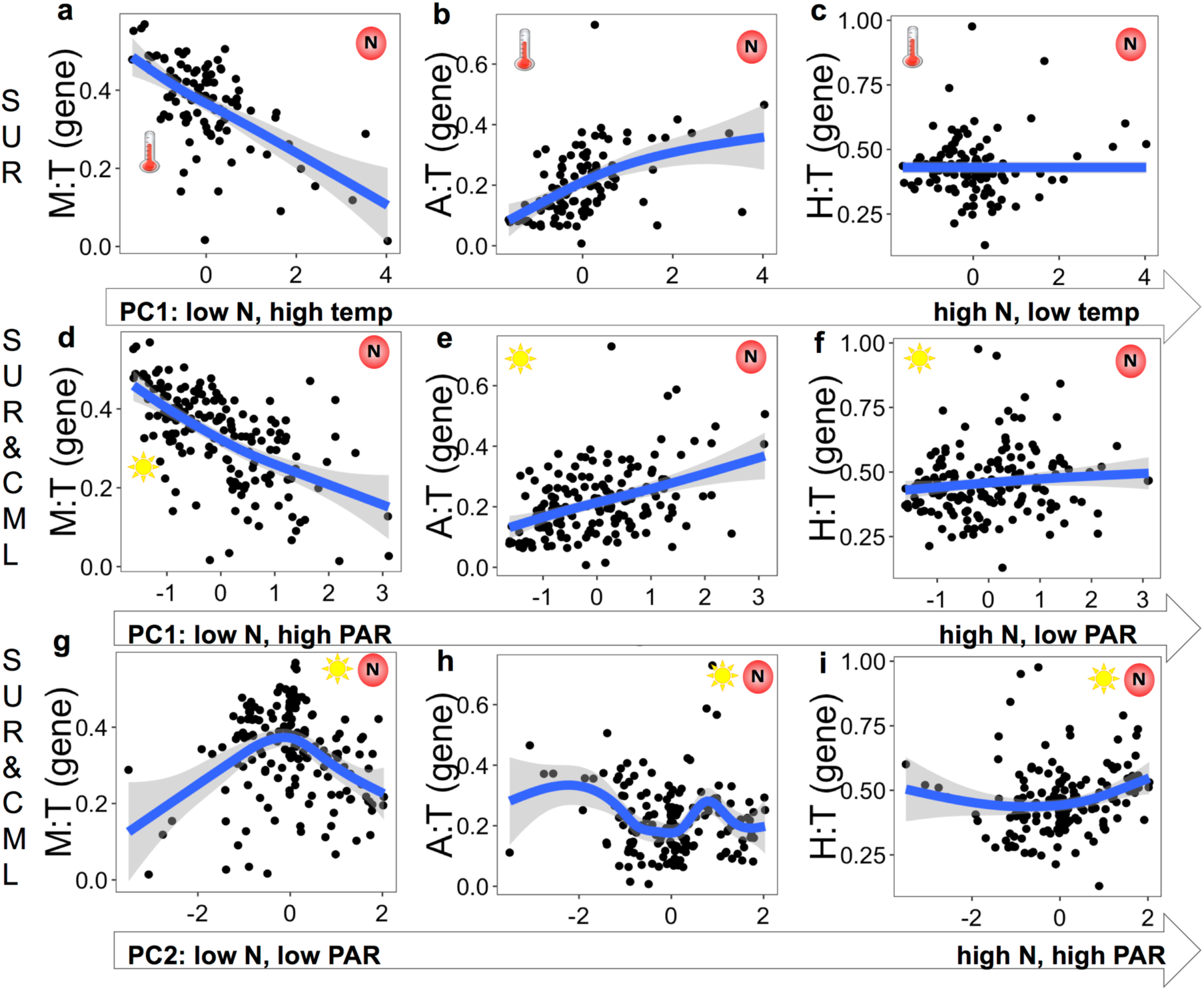
Variation patterns of TI_A/M/H_ in 18S rRNA gene abundances along PC1 axis for surface (SUR) dataset (**a-c**) and PC1 (**d-f**) and PC2 (**g-i**) axes for surface and chlorophyll maximal layer (SUR&CML) dataset. Principal components of nutrients (NO_3_^-^&NO_2_^-^), PAR, and temperature were marked on each panel, and the location of the symbols indicates higher values ends. Labels embedded on each PC-axis indicate how variables change from left to right. Blue lines and shaded bands were fitted generalized additive model smoother and 95% confidence interval.

Consistent with the PCA results, clear differences were observed for TI*_g_* between SUR and CML samples (from 61 stations have both SUR and CML samples), accompanied with distinct resources availability of nutrients, light and bacterial prey (**Fig. 4**). Significantly higher TI_M_ was seen in surface samples where PAR and bacteria were higher and N (NO_3_^-^&NO_2_^-^) were lower (median 0.29 vs 0.18, p<0.001, df=60). In comparison, TI_A_ was slightly higher in DCM compared to surface samples but not statistically significant (median 0.30 vs 0.29, p>0.05, df=60), and TI_H_ were significantly higher in CML than surface (median 0.52 vs 0.47, p<0.05, df=60) (**Fig. 4a, b**). Linear regression derived from all samples suggested a consistently positive relationship for TI_M_ with PAR and negative correlation with nutrients (NO_3_^-^&NO_2_^-^), from both SUR (122 samples; **Fig. 4c**) and SUR&CML dataset (165 samples; **Fig. 4d**). TI_A_ demonstrated a significantly positive and negative correlation with nutrients and PAR, whereas neither was significantly associated with TI_H_ (**Supplementary Figs. S2a-h**). For heterotrophic bacteria, a positive but weaker relationship (p=0.02) was found between TI_M_ and the SUR&CML dataset but not the SUR dataset (**Fig. 4c, d**). In contrast, both datasets have retrieved a strong negative association between heterotrophic bacteria and TI_H_, and a weaker but positive correlation between heterotrophic bacteria and TI_A_ in the surface (p<0.05) (**Supplementary Fig. S2i-l**). Other single variables showing significant regressions with TI_M_ were all shown in **Supplementary Fig. S3**.

**Fig. 4.**
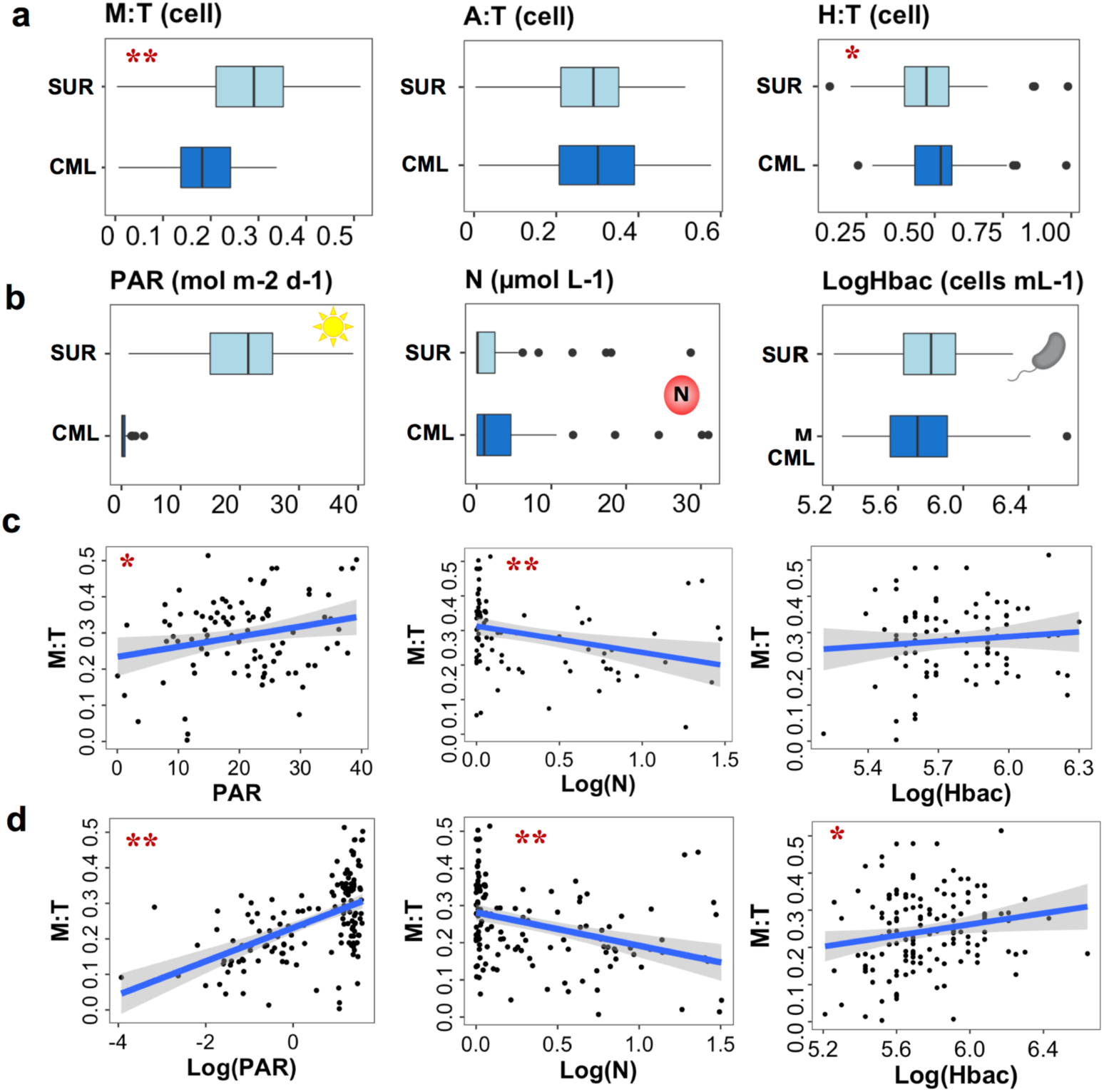
Comparison of TI*_g_*(**a**) and nutritional resource availability (**b**) between surface (SUR) and chlorophyll maximum layer (CML). Panels **c** and **d** demonstrate linear regressions between TI_M_ and three resource variables of PAR, NO_3_^-^&NO_2_^-^ (N), and heterotrophic bacteria (Hbac) with SUR (**c**) and SUR&CML dataset (**d**), respectively. Variables were sometimes Log-transformed for the best representation purpose. Significance codes were shown in red stars (one star if 0.001<P<0.05 and two stars if P<0.001), and shaded bands are pointwise 95% confidence interval on the fitted linear regression (blue lines).

### Mixotrophic growth with varying nutritional resource

The most abundant mixotrophs identified from the *Tara Oceans* dataset belonged to classes of Prymnesiophyceae, Dictyochophyceae, Chrysophyceae, Dinophyceae and Cryptophyceae (**Fig. 5a**). Mixotrophic strains we isolated from the North Pacific had been shown to be efficient grazers of heterotrophic bacteria and cyanobacteria (including *Prochlorococcus* and *Synechococcus*), and they were closely affiliated to those abundant *Tara Oceans* lineages [5,40]. Six strains selected for the experimental study were belonging to *Chrysochromulina*, Prymnesiophyceae_XXX of Prymnesiophyceae, Clade-H_X of Chrysophyceae, as well as *Florenciella*, Dictyochales_X and Florenciellales_X of Dictyochophyceae. Although the lineage from Florenciellales_X was not shown in Fig. 5a, it ranked as the 26th most abundant lineage across all stations. The relative abundances of those isolates retrieved from surface were all higher than CML except for Dictyochales_X (**Fig. 5b**), presenting similar patterns as the entire mixotrophic community (TI_M_ results in **Fig. 4a**). Given the consistent and substantial responses of TI_M_ to nutrients and PAR derived from field observations, we designed multifactorial bioassay experiments to assess impacts of light and nutrients (inorganic and organic nutrients) availability on the growth of mixotrophs (**Fig. 5c**).

**Fig. 5.**
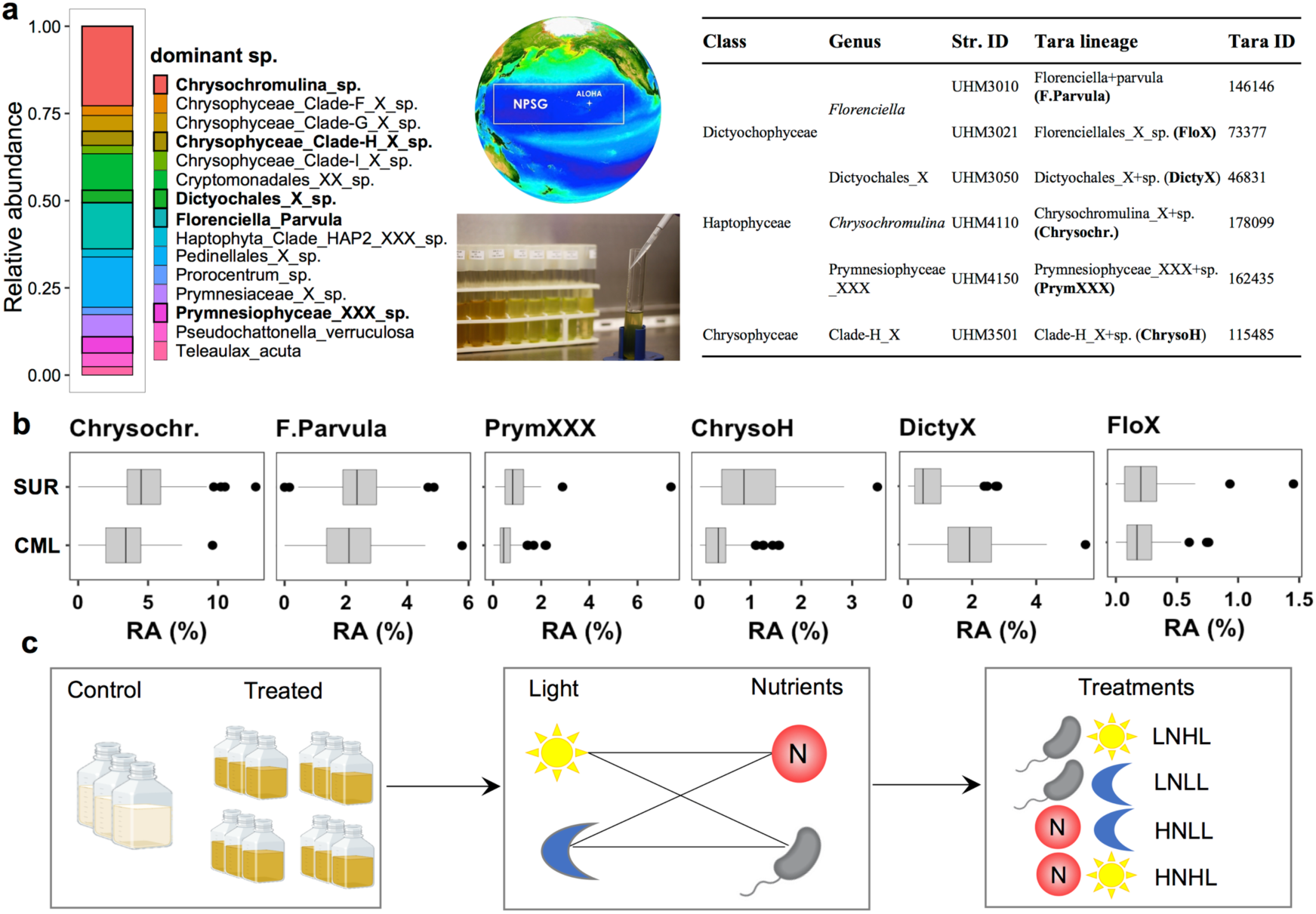
Relative abundances of top 15 most abundant *Tara Oceans* mixotrophic lineages (in gene converted cell abundances), and phylogenetically related strains isolated from North Pacific Subtropical Gyre (**a**). Comparison of relative abundances in surface (SUR) and chlorophyll maximum layer (CML) (**b**), and an illustrative flow chart showing how different treatments were set up for the growth experiments (**c**).

Distinct growth curve shapes and final abundances were presented among different treatments for the six isolates (**Fig. 6a**), accompanied by different disappearance rates of prey (**Fig. 6b**). Prey in controls (without mixotrophic grazers) all presented slower disappearance rates compared to cultures with grazers but to different levels among strains. Grazers in low nutrients control (without prey) grew to much lower concentrations, i.e., all <1×10^3^ cells mL^-1^ comparing to 10^4^-10^5^ cells mL^-1^ when added prey (data not shown). Shown as exponential growth rates (bars in **Fig. 6c**), most isolates presented the highest values when grown under high light, either when receiving high nutrients or sufficient prey (low nutrients). Lower growth rates were observed with low light treatments (marked with black arrows in **Fig. 6c**) for strain 4110 from *Chrysochromulina*, 4150 from Prymnesiophyceae_XXX and 3021 from Florenciellales_X, i.e., categorized as functional type I. The rest three isolates did not show significant impact by light, and thus were classified as functional type II. For grazing rates, most strains showed the highest clearance rates when receiving low nutrients and high light (marked with blue arrows in **Fig. 6d**). Only two strains belonging to Type II were capable of grazing much faster when receiving low light and high nutrients (marked with red arrows in **Fig. 6d**).

**Fig. 6.**
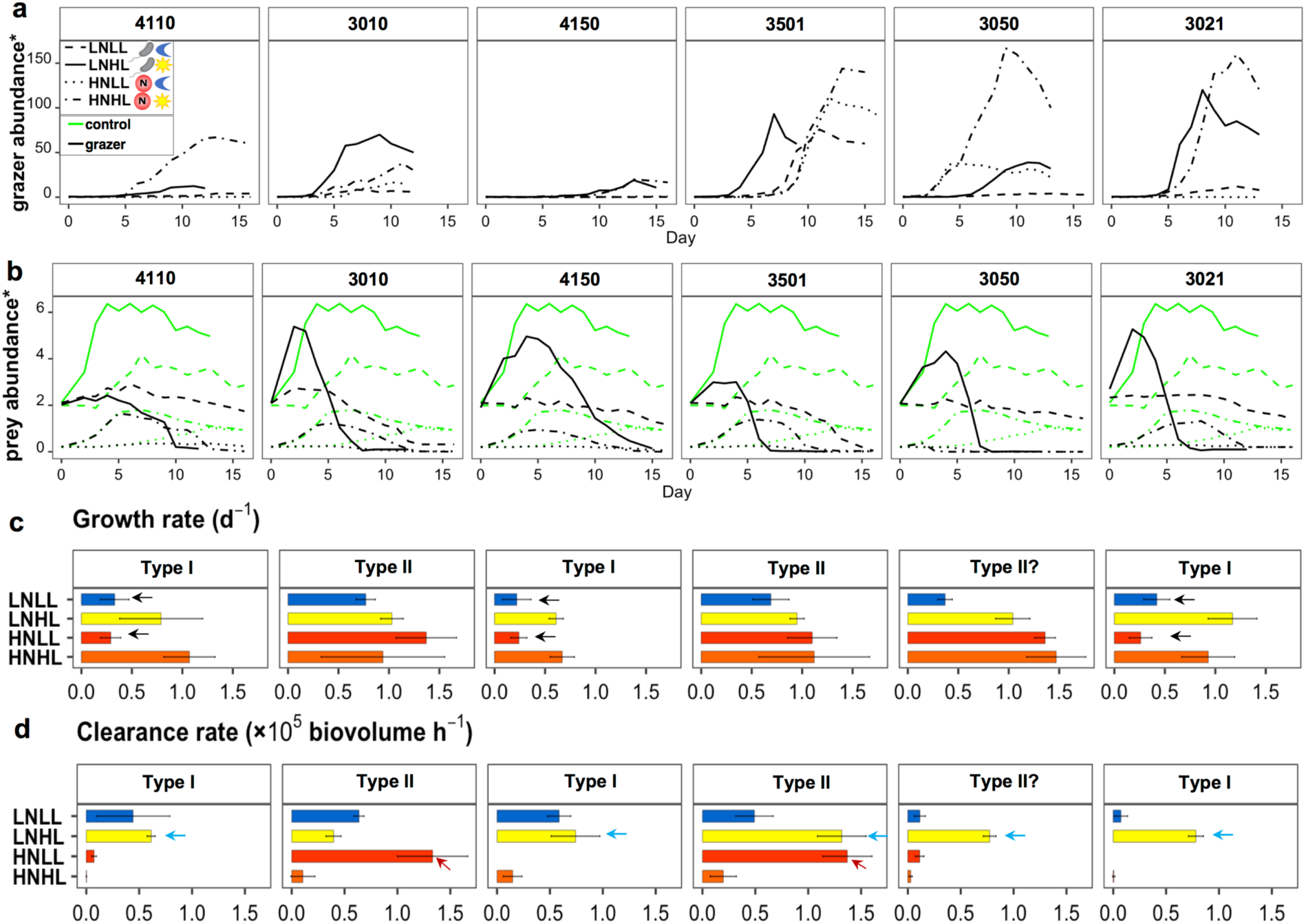
Panel **a** presents growth curves of six mixotrophic isolates grown under high light (HL) and low light (LL), as well as high inorganic nitrogen (HN) and low inorganic nitrogen (LN) conditions. All treatments were supplemented with *Procholorococcus* prey. Panel **b** presents removal of prey in all cultures (ID table in Fig. 5a); green lines indicate prey control without grazers and black lines were prey with grazers. Panels **c** and **d** demonstrate estimated growth rates during exponential phase and biovolume clearance rates for each strain. Errors bars in c and d denote standard deviation among replicates for each treatment but were not shown in panel a and b for more readability. **\***For grazer concentrations in HNHL, they were divided by 50000 cells mL^-1^ for #4110 and 5000 cells mL^-1^ for other strains, and 1000 cells mL^-1^ for all the rest treatments. *Prey abundances in all HN treatments were divided by 10^7^ cells mL^-1^, and all LN were divided by 10^6^ cells mL^-1^. Arrows in **c** and **d** indicate treatments that had caused significantly lower growth rates (c), and significantly higher grazing rates (d).

ANOVA results revealed a significant effect on growth by light, i.e., to increase growth rates with increasing light, either alone, or together with nutrients and species. However, (inorganic) nutrients availability did not show significant impact on growth rates, as cultures supplemented with bacterial prey can sufficiently compensate for nutrients deficiency. Less significant effects were observed for clearance rates, as strains responded differently to changing light and nutrients conditions, indicated by the high p (>0.05) values from both single and multi-factorial ANOVA (**Table 3**).

**Table 3.**
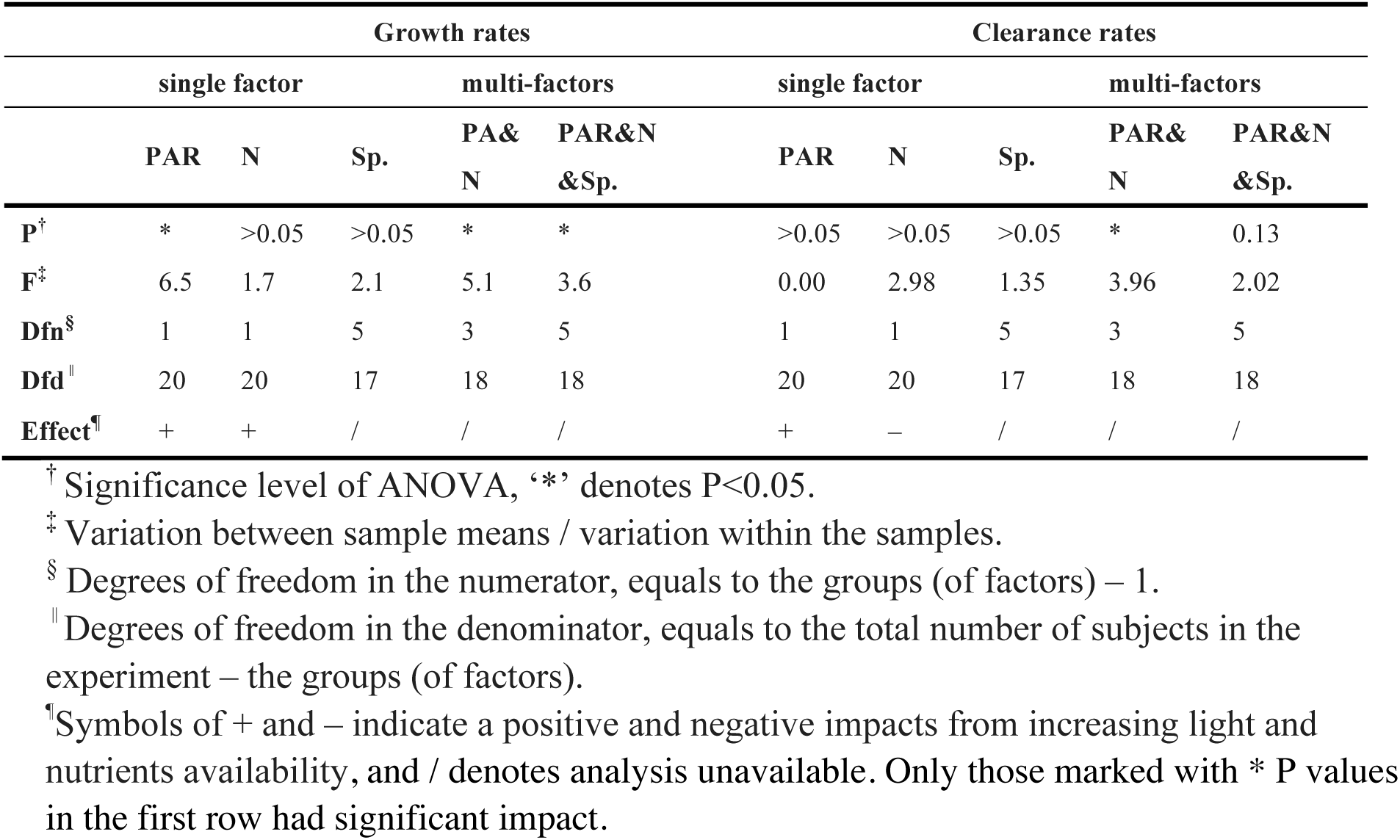
ANOVA of different growth rates and clearance rates among six species cultivated under different light (PAR) and nutrients (N) conditions (results shown in Fig. 6). Individual and interactive effects of light, nutrients and species were analyzed via single factor and multi-factors ANOVA, respectively.

## Discussion

In this study, we used large-scale field surveys to retrieve prominent variables that possibly affect the trophic mode structure of eukaryotes on community levels, with a particular focus on three nutritional resources. We then conducted finely controlled lab experiments using isolates to investigate how mixotrophic growth respond to changing nutritional resources, in terms of both forms (inorganic nutrients and organic nutrients) and availability. While acknowledging species-specific variations, both field and experimental results retrieved a close relationship between mixotrophic prevalence and nutritional resource among inorganic nutrients, light and prey, which have possibly introduced distinct niche preferences for mixotrophs versus autotrophic and heterotrophic eukaryotes in the ocean.

### Mixotrophic growth responding to changing nutritional resources

Suggested by our experimental results, mixotrophy can function as an efficient strategy for compensating inorganic nutrients deficiency for most strains through ingestion of bacterial prey, but not necessarily for supplementing light limitation (Fig. **6c**). Among the six studied cultures, three of them were efficient with compensating light/energy deficiency via heterotrophic nutrition (grazing), but not the other three. These results indicated that mixotrophic controlling mechanisms for metabolic partitioning between autotrophy and heterotrophy were species-specific. The Type I isolates that were light-dependent possibly align with the ‘primarily photosynthesis and phagocytosis for nutrients’ functional group proposed by Jones (1997) and Stoecker (1998) [51] [52]. The other three strains of Type II were light-independent, and likely belong to the ‘phagocytose when light is limiting’ or ‘perfectly balanced mixotrophy’ functional group. Based on proposed resource allocation theories, those light-dependent mixotrophs probably rely on photosynthesis for gaining most energy and organic carbon [12,13,14]. The light-independent group performed better with low light availability, suggesting a higher flexibility governing autotrophic and heterotrophic growth in response to the changing conditions.

### Relationship between mixotrophy and nutritional resources on large spatial scale

Our field analysis also indicated that availability of three nutritional resources could affect the relative success among three trophic strategies on community level, and across large spatial scales. Specifically, higher-nutrients and lower-light environments could possibly promote the relative abundances of autotrophs, whereas mixotrophic community proliferated from low nutrients and high light environments. This positive light-mixotrophy relationship in the field can be partially explained by the possibility that the light-depend Type I mixotrophs were dominant across the investigated ocean regions. On the other hands, Rothhaupt (1996) and Edwards et al. (2019; 2022) have suggested that during a resource competition process, mixotrophs could outcompete heterotrophs, because they can use extra energy gained from photosynthesis which enable them to suppress bacterial prey to very low level (below the threshold of heterotrophic grazers) [8,11,53]. Both mechanisms could be true, and it is worthy to conduct further studies to investigate the fundamental mechanisms related to metabolic partitioning and resource allocation for mixotrophic strategy.

Heterotrophic bacteria maintained relatively consistent abundances in the sampled oceans and was not selected as main explanatory factor for TI*_g_* by RDA or PCA. Nevertheless, they demonstrated as one of the predictors explaining TI_M_ based on stepwise regression model (**Supplementary Table S2**). When treated as a single variable, a strong negative correlation was found between TI_H_ and heterotrophic bacteria. Comparatively, the correlation was positive for TI_A_ and less significant for TI_M_. This interesting shifts likely suggested distinct interactions between bacteria and three trophic groups. Heterotrophic bacteria benefit from the dissolved organic carbon (DOC) excreted by autotrophs and recycle nutrients for autotrophic growth, forming a positive relationship. Whereas for heterotrophic eukaryotes, this negative association could be attributed to high grazing pressure they exerted on heterotrophic bacteria [43,44]. The weaker but positive correlation between heterotrophic bacteria and TI_M_ could be explained by the more complex relations among bacteria (prey and decomposer) and mixotrophs (grazer and DOC provider), as well as the lower grazing capability/pressure from mixotrophic grazers than heterotrophs [13,23,38,45].

### Environmental variables could affect the prevalence of trophic strategies

Other oceanographic conditions such as CML, nitracline depths and MLD also seemed to affect mixotrophic prevalence, though they are physical processes that are more or less associated with nutrients and light conditions. For example, deeper nitraclines can potentially suppress nutrients supply in the surface and therefore increase TI_M._ Similarly, deeper chlorophyll maxima could be indicative of higher light availability and lower nutrients in photic depths, thereby enhancing TI_M_. In contrast, deeper mixed layer depths (stronger mixing) could reduce light availability and increase nutrients concentration, which both weaken TI_M_. The overall positive association between CML and nitracline depths and TI_M_, and negative correlation between MLD and TI_M_ were in line with their overall relationships with nutrients and light.

Previous field studies suggested that besides nutrients [21,46] and light [47,48,49], variables such as temperature [50] and carbon composition (colored DOM and CO_2_) [20] were also likely important for affecting mixotrophic prevalence. These variables were also included in our analysis and yielded varying relationships with TI*_g_*. These differences could possibly be due to the different environmental gradients covered among our studies (various open oceans) versus others (North Pacific, and boreal Canadian lakes). Stepwise regression model suggested that PAR, nutrients, density, PIC, DCM, heterotrophic bacteria and *Prochlorococcus* were the top predictors explaining TI_M_ with the *Tara Oceans* dataset (**Supplementary Table S2**).

Indeed, it is not always straightforward to delineate relationships between a single environmental variable and trophic mode prevalence. Results from our statistical analysis rather provide insights into the possible correlations between them. Future studies focusing on smaller regions with in situ experiments targeting active trophic behaviors will provide additional knowledge for interpreting the trophic mode and environment relationships.

### Methodology caveats

One potential caveat in our methodology is that mixotrophs were annotated based on published literature at species or higher taxonomic ranks. These trophic annotations were not necessarily aligning with in situ metabolic activities. Nevertheless, the potential or historically evolved capability of phago-mixotrophy are sufficient to delineate their ecological niche which was the focus of this study. We also recruited longer-term averaged values for some of the key variables (e.g., 8-day and 30-day PAR) as well as a variety of parameters (e.g., NO_3_^-^, NO_2_^-^, PO_4_^3-^ and SiO_4_^-^ for nutrients) to reduce the possibility of retrieving false relationships. Lack of trophic evidence for some of the species/lineages could on the other hand, cause false annotation, and variations against other studies. For example, we annotated *Micromonas*, a member of Chlorophyta, as autotrophs based on the best knowledge of ours [51,54]. Howerver, others have argued possible phagotrophic evidence in the *Micromonas polaris* [55]. We also recognize the gene-cell abundance correction factors could be inaccurate for certain groups, as we did not have first-hand cellular abundance data from in situ samples. Nevertheless, these calibrated abundance data could improve accuracy for some heavily over-estimated groups such as dinoflagellates and under-estimated small flagellates.

Combining various approaches including experimental study, field observation, model simulation and bioinformatics tools can be powerful to retrieve the most reliable data for future mixotrophy research [3,56].

## Conclusion

This study provides the first *Tara Oceans* analysis for niche partitioning among three trophic populations of mixotrophs, autotrophs and heterotrophs of the eukaryotic community. Our results suggested that either on community level or species level, phagotrophic mixotrophy are a trophic strategy marine eukaryotes had evolved to survive nutrients impoverishment and light limitation, deepening our understanding about mechanisms controlling their prevalence in the ocean.

## Acknowledgements

We thank the *Tara Oceans* project and their team for providing the original database of eukaryotic 18S rRNA metabarcodes and environmental parameters. We also acknowledge the funding resource of Young Scientists Fund of the National Natural Science Foundation of China, and the Oceanic Interdisciplinary Program of Shanghai Jiao Tong University.

---

**Supplementary Table S1.**
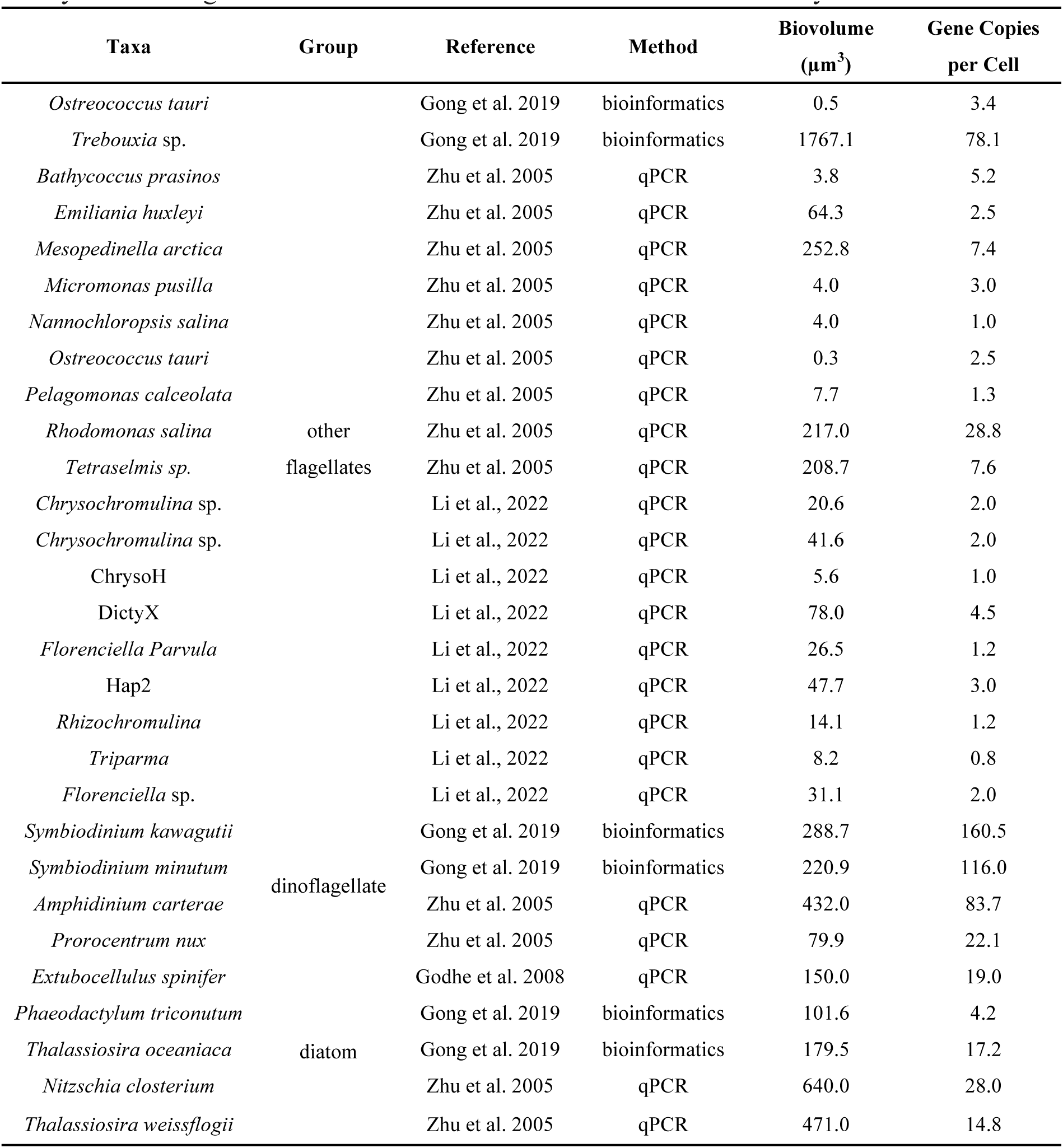
Species and cellular 18S rRNA gene copy numbers used in this study to retrieve gene-cell abundance correction factors for small eukaryotes.

**Supplementary Table S2.** Stepwise regression analysis revealed significant variables contributing to the prevalence of mixotrophs in both surface and chlorophyll maximal layer (SUR&CML) samples, and surface (SUR) samples alone. Significance of P values and coefficient for 8 selected predictors were given in each row.

| SUR&CML |  |  | SUR |  |  |
| --- | --- | --- | --- | --- | --- |
| Predictor<sup>†</sup> | P<sup>‡</sup> | Coefficient | Predictor<sup>†</sup> | P<sup>‡</sup> | Coefficient |
| PAR | ** | 0.1 | CML | *** | 0.15 |
| NO<sub>3</sub><sup>-</sup>&NO<sub>2</sub><sup>-</sup> | *** | -0.08 | *Prochlorococcus* | ** | -0.82 |
| Density | *** | 0.80 | NO<sub>3</sub><sup>-</sup>&NO<sub>2</sub><sup>-</sup> | *** | -0.15 |
| PIC | ** | 207.5 | PIC | * | 218.3 |
| CML | *** | 0.10 | PO<sub>4</sub><sup>3-</sup> | * | 0.28 |
| Hbac | ** | 1.25 | Density | * | 0.78 |
| *Prochlorococcus* | * | -0.53 | PAR | ** | 0.12 |
| / | / | / | NPP | . | 0.08 |

<sup>†</sup>Abbreviations: PAR, photosynthetically available radiation; PIC, particulate inorganic carbon; Hbac, heterotrophic bacteria; NPP, net primary production
<sup>‡</sup>Significant codes: '\*\*\*' p<0.001, '\*\*' p<0.01, '\*' p<0.05, '.' p<0.1

## Supplementary figures

**Fig. S1.**
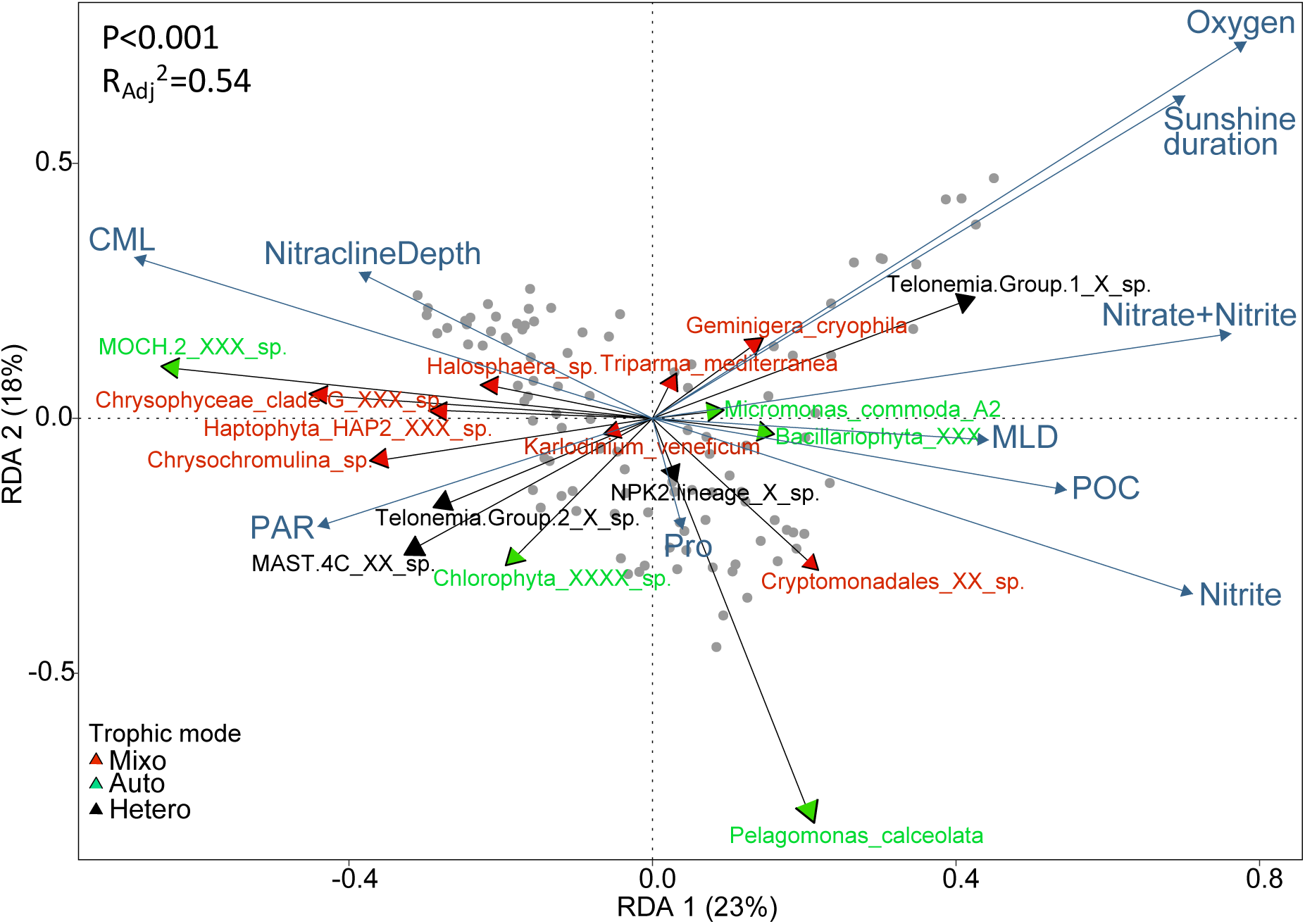
Redundancy analysis for environmental impact on the distribution of trophic groups in surface oceans, derived from 18S rRNA gene corrected cell abundances. Only 17 *Tara Oceans* lineages passed Escoufier selectin (0.9 threshold) were shown, and adjusted R^2^ of the analysis is of 0.54. Environmental variables selected were denoted in cyan-blue arrows and text. Abbreviations: PAR, photosynthetically active radiations; CML, chlorophyll maximum layer; MLD, mixed layer depth; POC, particulate organic carbon; PIC, particulate inorganic carbon; Pro, *Prochlorococcus*.

**Fig. S2.**
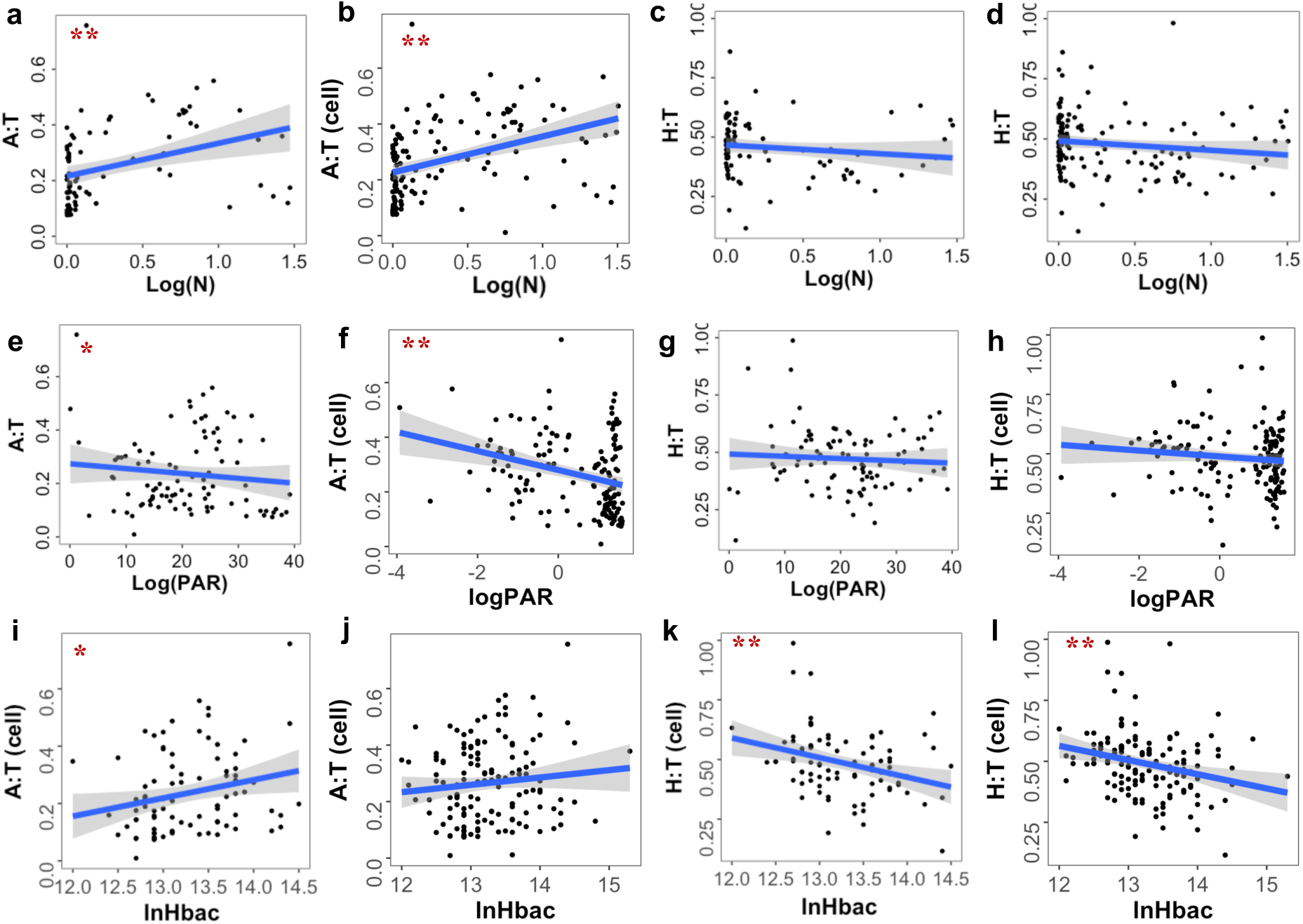
Linear regression between TI_A/H_ (all in cell abundances) and resource variables retrieved from the SUR (104 samples) and SUR&CML dataset (165 samples). Different significance levels were marked with different number of red stars (one star if 0.001<p<0.05 and two stars if p<0.001), and shaded bands are pointwise 95% confidence interval on the fitted values (the line). Abbreviations: PAR, photosynthetically active radiations; N: inorganic nutrients of nitrate (NO_3_^-^) and nitrite (NO_2_^-^); Hbac: heterotrophic bacteria.

**Fig. S3.**
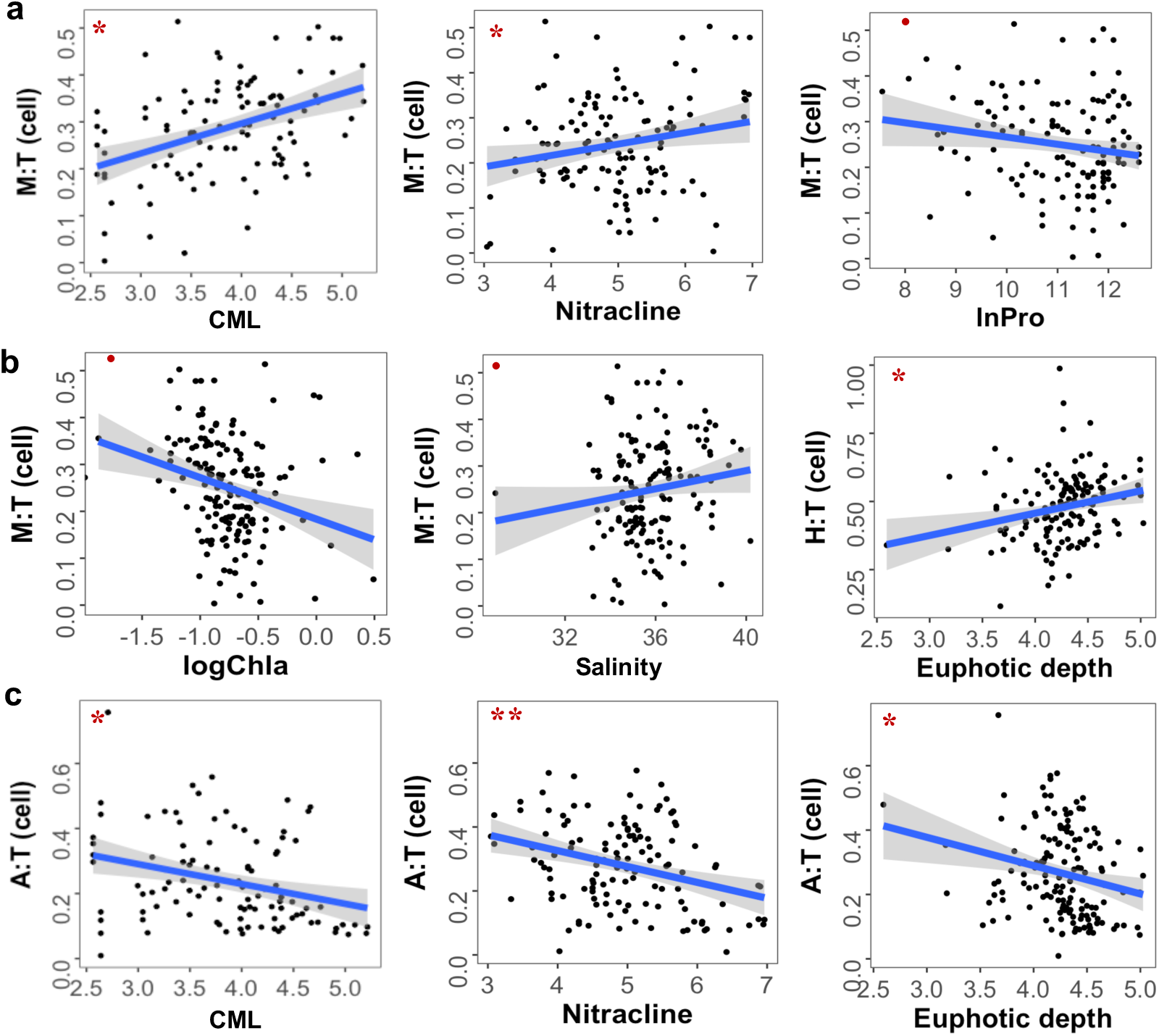
Linear regression between TI*_g_* (all in cell abundances) and single environmental variables that have shown significant correlations, retrieved from all 165 SUR&CML samples. Different significance levels were marked with different number of red stars (one star if 0.001<p<0.05 and two stars if p<0.001), and shaded bands are pointwise 95% confidence interval on the fitted values (the line). Abbreviations: CML, chlorophyll maximal layer; Chl a, total chlorophyll a, Pro, *Prochlorococcus*.

